# Evaluating the Spatial Overlap of Cattle and Wildlife on Buck Island Ranch, Florida

**DOI:** 10.64898/2026.08.14.744666

**Authors:** Sulagna Chakraborty, Olivier Massicot, Brian F. Allan

## Abstract

Spatial overlap of domestic animals and wildlife can have positive effects such as increased biodiversity in some areas and reduction in forest fires but it can also have negative consequences such as pathogen spillover, vector and pathogen propagation into new areas among others. Thus, understanding the interactions between wildlife and domestic animals on shared landscapes is critical for conservation and management purposes. To investigate these cattle-wildlife interactions, we chose Buck Island Ranch (BIR) in southcentral Florida. We studied spatial overlap between Domestic Cattle (*Bos Taurus*) and multiple wildlife species such as Wild Boar (*Sus scrofa*), White-tailed Deer (*Odocoileus virginianus*), Rabbits (*Sylvilagus* spp) and others in pasture, semi-pasture, and hammock habitats/vegetation types at BIR. Five management blocks were selected that had pasture and hammock habitats and within each block we established matched pairs of 100m transects. Animal dung samples were surveyed and identified in 13 pairs of transects. We used Wilcoxon signed rank test to determine differences in dung counts between the habitats and calculated Simpson diversity index to analyze the differences in diversity between the habitat types. We found that there was greater dung count of white-tailed deer in hammock habitats compared to pastures (p=0.048) and greater diversity of wildlife in hammocks than pastures (p=0.028). Cattle dung was found in high density in both hammocks and pastures. These results indicate that hammocks are areas shared between cattle and wildlife at BIR and depending on the extent of direct and indirect contacts between these animals there can be a risk for pathogen spillover. We recommend further research on local land and cattle management practices that can facilitate the co-existence of livestock and wildlife.

## Introduction

The livestock industry is a large economic sector in the United States where the highest value of production comes from the Domestic Cattle (*Bos taurus*) industry estimated at $50.2 billion in 2017 (USDA APHIS 2017). Livestock are essential as a source of income (FAO 2023), are important in reducing food insecurity, drive economic growth, and empower people and pastoralists throughout the world who rely on them culturally and fiscally (Banda and Tanganyika 2021). However, livestock ranching also pose negative consequences to the environment. The livestock sector is an important user of natural resources and has considerable influence on air quality, global climate, soil quality, biodiversity, and water quality (Tullo et al. 2019). With the increase in livestock production globally, there is increasing spatial overlap and competition between wildlife and livestock for available land, food, habitats, and other resources, which can have negative consequences on biodiversity (Gordon 2018). These interactions in turn can lead to undesirable ecological changes including opportunities for pathogen spillover between wildlife and livestock (Karmacharya et al 2024). The grazing effect of cattle can also impact the wildlife in these areas, for instance, in a review of literature focusing on cattle-deer interactions in North-American ecosystems, it was found that compatibility between cattle and deer were driven by geographic region, cattle stocking rates, and season (Hines et al 2021). Thus, to prevent the loss of wildlife diversity and negative environmental impacts, sustainable methods of agriculture and management scenarios need to be implemented which allow both wildlife and livestock to cohabitate. A growing body of research indicates possible benefits of having wildlife and domestic animals coexist (Jori et al. 2021), such as adoption of land management practices closer to the local natural conditions (Pywell et al. 2012), aiding in conservation of wildlife and increasing biodiversity (Allan et al. 2017), and increasing revenue from wildlife tourism (Melita et al. 2013). For these reasons, investigation of real-world management scenarios where there is regular interaction or spatial overlap between wild and domestic animals is a crucial step to address possible harmful or beneficial effects from livestock production.

Surveys to estimate abundance of wildlife species, and to a lesser extent, domestic livestock, can present logistical challenges and expenses. Dung sampling is a cost-effective method used to estimate density or activity levels of animals in a given area and can also be used to determine trends in animal populations over time (Laing et al 2003). Dung sampling might not provide precise information such as age of animals or the population size (Mohanarangan et al 2022; Hedges et al 2013); additionally factors such as defecation rates of the animal, period of dung sampling, and dung decay rates need to be accounted for while making abundance estimates (Laing et al 2003). Dung sampling is useful when methods such as aerial or mark-recapture surveys are not possible or in areas where direct monitoring of animals is difficult without disturbing them. Biologists have used dung counts for surveying wildlife such as African Elephants *(Loxodonta*) and Asian Elephants (*Elephas maximus)* (Barnes 2008), Snowshoe Hares (*Lepus americanus)* in southwestern Yukon (Krebs et al. 2001), White-tailed Deer (*Odocoileus virginianus)* in Mexico (Camargo-Sanabria and Mandujano 2011), Gorillas *(Gorilla gorilla*) in central African Republic (Todd et al. 2008), and many others. Several studies have used this method to determine the ecological processes that occur in areas where cattle and wildlife use shared habitats such as among cattle and wild herbivores in Laikipia, Kenya (Keesing et al. 2018), among cattle and Red Deer (*Cervus elaphus*) in alpine pastures in Italy (Forti et al. 2021), niche breadth of Swamp Deer (*Rucervus duvaucelii*) in comparison to Spotted Deer (*Axis axis*) and domestic cattle in Nepal (Regmi et al. 2022), and Nesting Greater Sage-grouse (*Centrocercus urophasianus*) and cattle in Montana (Smith et al. 2018).

To understand the degree of spatial overlap between livestock and wildlife, we used livestock and wildlife dung sampling in a specific cattle-wildlife integrated system in North America. The objectives of this study were to determine i) differences in the dung counts of cattle and wildlife by habitat/vegetation type and ii) differences in species diversity by habitat/vegetation type. The results of this study will help unravel areas that are shared by these different animals and potential implications for the same.

### Field-Site Description

Buck Island Ranch (BIR) that is associated with Archbold Biological Station in Venus, Florida, is one such site in North America where the goal is to have active cattle production that is compatible with wildlife. In addition, BIR has been the subject of numerous ecological studies, providing additional context to understanding wildlife-livestock interactions. Since only few such studies have been conducted in North America, BIR is an ideal setting to investigate these interactions and contribute to the current understanding of how spatial overlap of wild and domestic animals occurs under the conditions of an operating cattle ranch.

Buck Island Ranch in Venus, Florida is a 4,249-hectare cattle ranch supporting an estimated ∼ 3,000 head of cattle at the time of this research. BIR has two pasture systems: improved, and semi-native grassland, which is further divided into numerous management blocks (Sonnier et al. 2020, <u>Archbold Biological Station/habitats</u>). Ranch managers rotate herds of cattle among the pastures and management blocks according to seasonal and environmental management objectives (<u>Archbold Biological Station/cattle</u>). The improved pastures are dominated by Bahia Grass (*Paspalum notatum)* whereas the semi-native ones (also referred to as “unimproved”) are dominated by Field Paspalum (*Paspalum dilatatum*), bunch grasses such as Broomsedge *(Andropogon virginicus* L.), Bushy Bluestem *(Andropogon glomeratus*), and Panic Grass (*Panicum longifolium*), along with a variety of Sedges (*Cyperaceae* spp.) (Archbold Biological Station; n.d.). The two types of pasture also differ in that the improved pastures have been historically ditched, fertilized, replanted with Bahia, and are typically at a higher elevation making them drier during the wet season. The semi-native pastures have not been fertilized or extensively ditched, have a greater percentage of native species, and are at a lower elevation resulting in inundation during the wet season (Archbold Biological Station; n.d.).

In both improved and semi-native pastures there are naturally occurring hammocks, which are areas of dense tree cover consisting of Cabbage Palm (*Sabal palmetto*), Live Oak *(Quercus* spp.), and Spanish Moss (*Tillandsia usneoides*), with an understory of open shrub layer, sparse herb layer and senescent leaf litter (Merrill et al. 2018). Hammocks are interspersed throughout the pastures on BIR and likely provide important habitats for wildlife, including white-tailed deer and Feral Pigs *(Sus scrofa*), and shade for cattle (<u>Archbold Biological Station/habitats</u>, Merrill et al. 2018). The soil in BIR is primarily composed of fine sand hyperthermic alfisols and spodosols with very poor nutrients (Kohmann et al 2021), while the climate in BIR is warm dry season from November to May, and a hot wet season from June to October, with 75% of the rainfall in the wet season (Qiu et al 2024). Land managers at Archbold Biological Station use a prescribed fire management program to restore near-natural fire regimes and provide a template for studying the ecology of natural and altered fire regimes (<u>Archbold Biological Station/Prescribed Burns</u>)

## Methods

To assess the extent of spatial overlap between cattle and wildlife, we sampled both types of pastures in comparison to hammock habitat/vegetation types for the presence and density of dung from cattle, other domestic animals, feral pigs, and all detectable species of wildlife. Based on the configuration of management blocks on BIR, we selected five management blocks wherein we could sample both pastures and hammocks. Blocks were sampled when cattle were not present to avoid disturbance. Within each block, we established matched pairs of 100m transects in both vegetation/habitat types (i.e., hammock transects paired with pasture transects). Transect locations were selected at random to maintain a minimum distance of 90 meters from other transect pairs and 40 meters from roads and trails but were otherwise placed randomly in every block. Adhering to these criteria we were able to establish 1 to 4 pairs of transects per block, according to the area of the management block and naturally occurring hammocks within those, thus we had a total of 26 transects (13 pairs) in which we conducted dung sampling. In some instances, randomly placed transects intersected with supplemental feed plots for livestock. In all the selected transects, dung sampling was performed in the same manner as further described. Additionally, transects were also selected to ensure that they did not traverse water bodies or did not fall in management blocks where cattle might be present. Given these constraints, we were only able to sample in one block that had the improved pasture type, most of the pasture habitats sampled were of the semi-native type. Table 1 provides a description of the management blocks, number of transects selected in each block, and information provided by property managers on time since last cattle grazing and time since last prescribed fire.

**Table 1.**
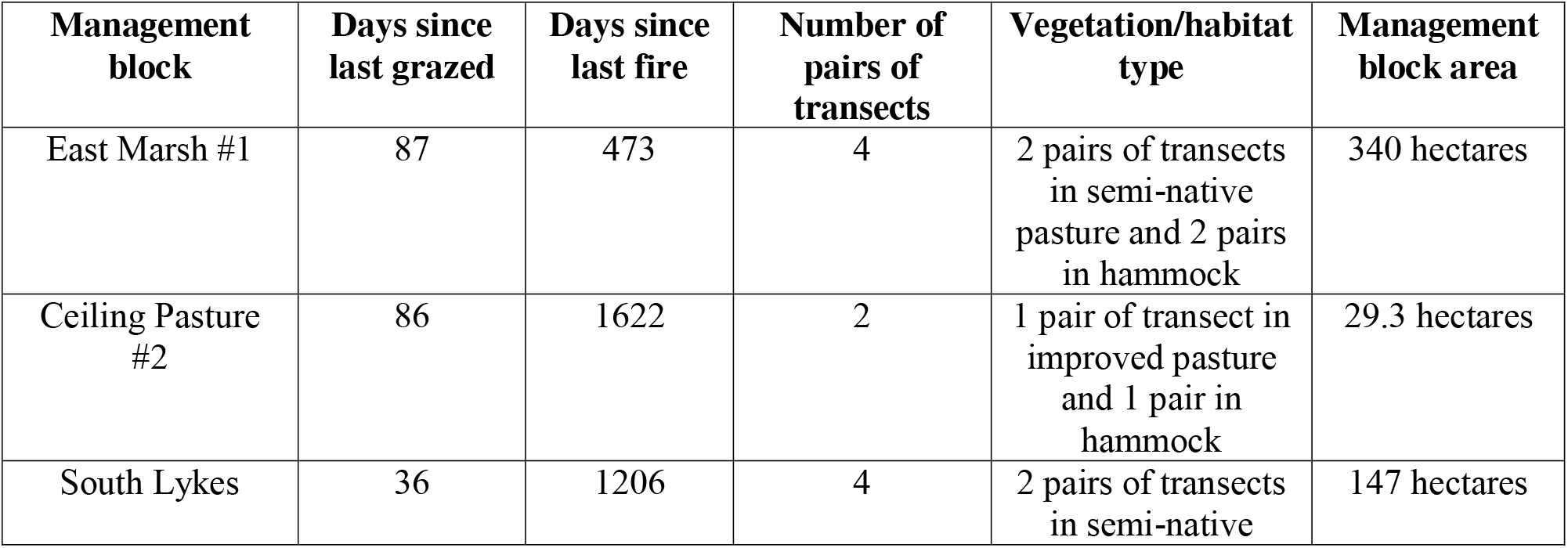
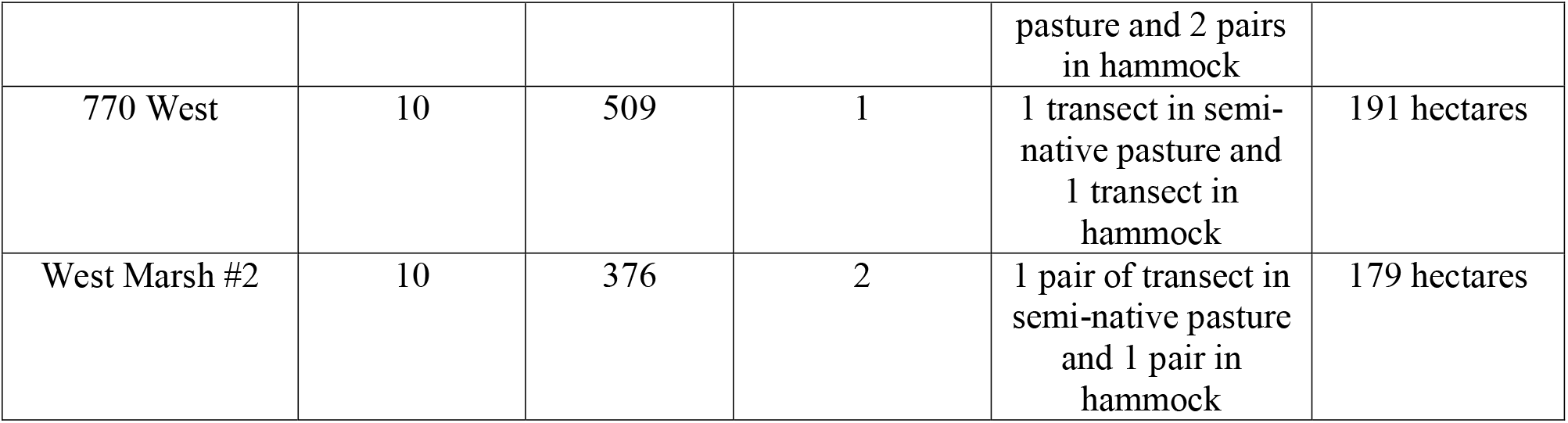
Dung survey sites at Buck Island Ranch, Archbold Biological Station, Venus, FL.

We conducted the field sampling for this project from 25 May 2021 until 5 June 2021. Sampling was performed from morning hours to early afternoon, typically between 7:30 am and 1:00 pm. We counted and identified all detectable animal scat (Table 2) to up to 2m on either side of the 100m transects (400m^2^ sampled per transect) and collected the GPS coordinates for the start and end points for each transect using a Garmin Rhino GPS (Garmin, Lenexa, Kansas) (Fig 1).

**Table 2.**
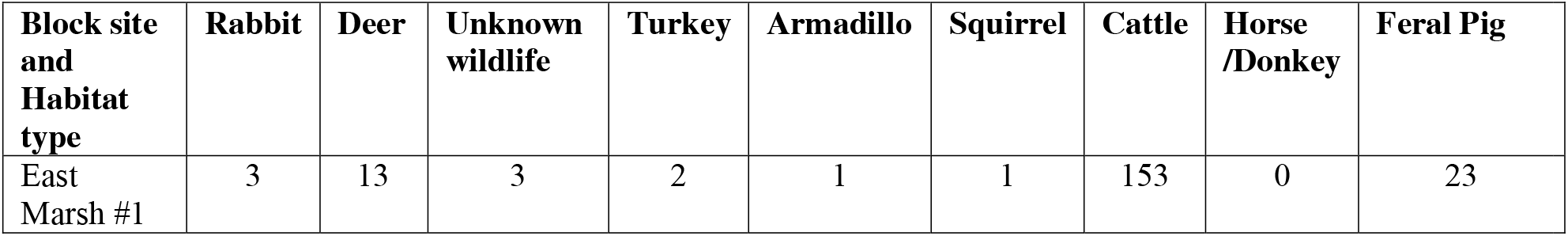
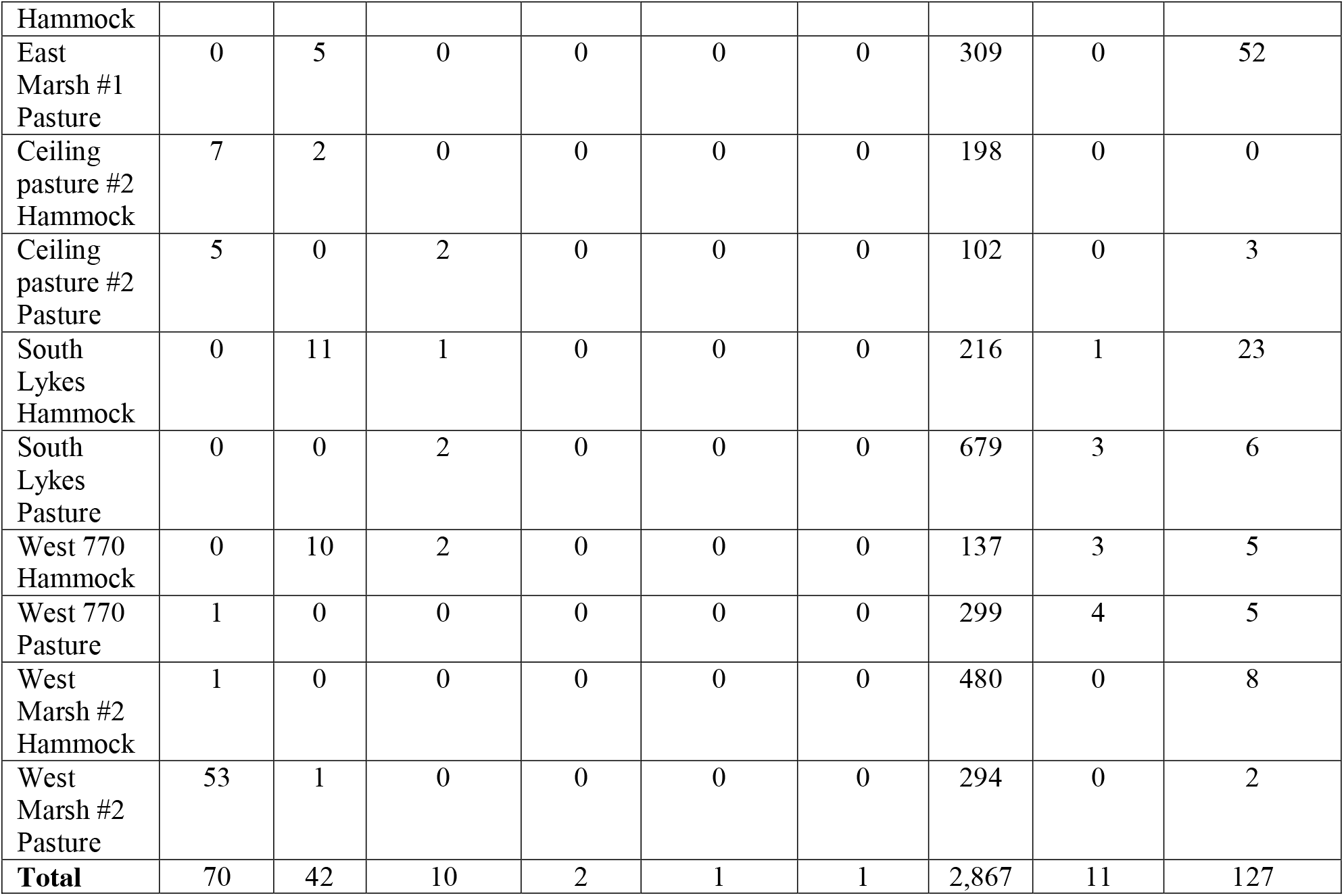
Dung counts of all animals detected in each of the sampled sites (i.e. 13 pairs of transects in 5 blocks) and in all the habitats/vegetation type (pasture vs hammock) at Buck Island Ranch, Archbold Biological Station, Florida. *Note: here we provide the sum of all dung counts of detected animals across all transects within each block*.

**Figure 1.**
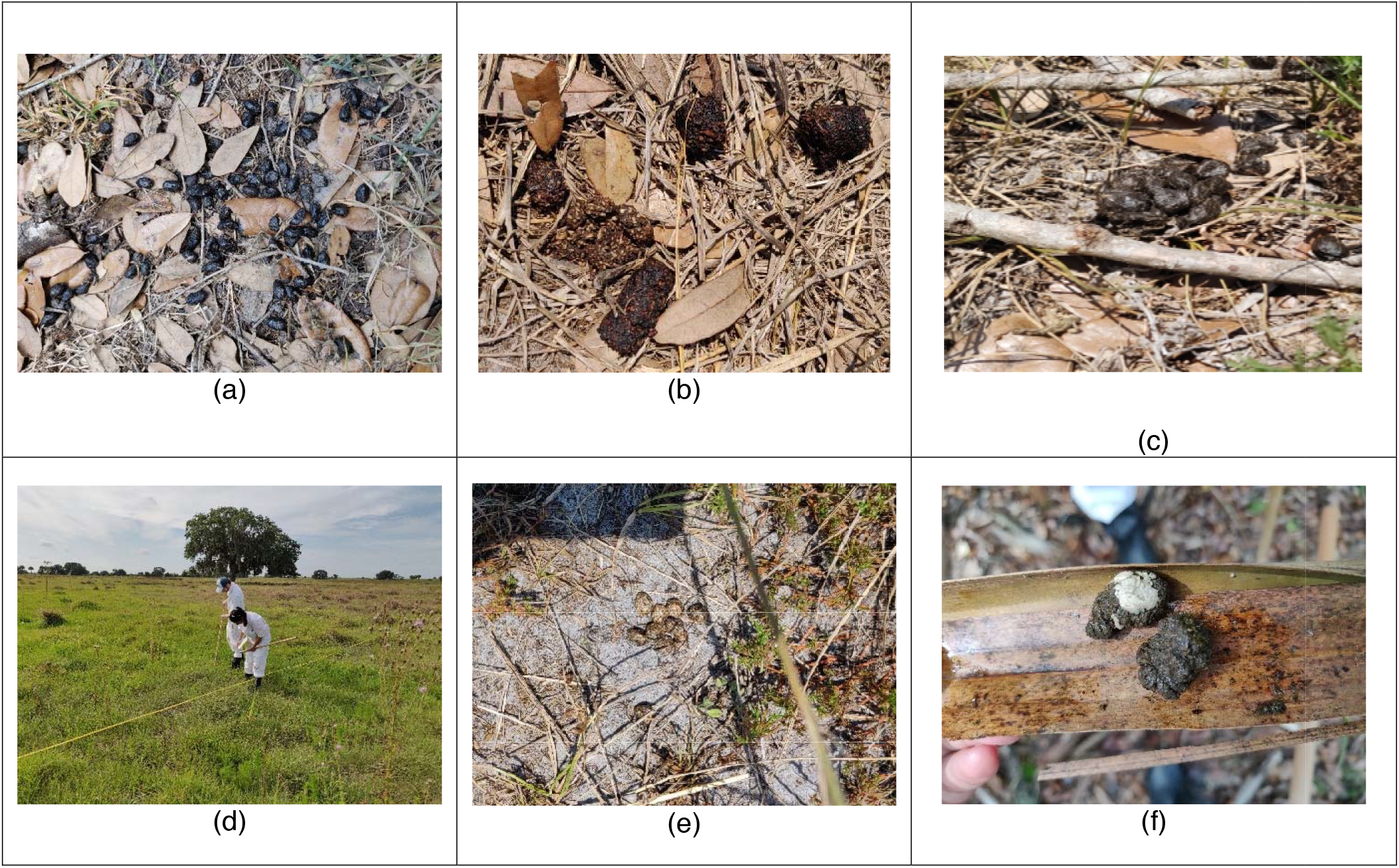
From top left to right (a) a cluster of Deer dung; (b) a meso-carnivore dung sample; (c) a cluster of Feral Pig dung (d) a transect in pasture habitat that was comprised of a mixture of Bahia and improved grass; (e) a cluster of Rabbit dung, and (f) a Wild Turkey scat sample. ***Note***: *In photograph 1d, the authors of this paper (SC, OM) are sampling in a pasture habitat, picture was taken by the third author (BFA)*.

Two observers conducted the dung counts in each transect and took photographs of any unidentifiable scats. We utilized expert opinion and guides for animal sign and scat (Elbroch and McFarland 2019) to aid in the identification of dung. Photographs of unidentified samples were also uploaded as part of this project on the crowdsourcing platform iNaturalist (https://www.inaturalist.org/) to gain assistance in identifying unknown dung samples. IACUC approval was not required for this research since we did not handle or interact with live animals.

Statistical analyses of dung counts observed along each transect of hammock and pasture vegetation were performed in R v.4.2.0 via R Studio (Posit team 2025). Because the data were not normally distributed, we conducted Wilcoxon signed rank tests to determine whether there was a difference in the dung counts in hammock versus pasture habitats. We performed a square root transformation of the dung counts to have a symmetric distribution of the data, then repeated the Wilcoxon signed rank tests to see if there was a difference in dung counts between hammock and pasture habitats. Because some species were observed rarely, statistical analyses were only performed for species with a minimum of 12 observations across all transects (i.e., cattle, feral pigs, white-tailed deer, and all wildlife species combined). Paired t-tests were run following which we ran Wilcoxon signed rank test with continuity correction for those variables as described above. We calculated effect sizes between cattle and feral pigs dung samples and the combined dung samples of cattle and feral pigs with that of wildlife for comparison in Microsoft Excel (Microsoft Corporation 2024). Additionally, we calculated the Simpson’s diversity index (Simpson 1949) of dung samples for all animal species and for only wildlife in order to analyze the differences in diversity between hammock and pasture habitats. We used Welch Two Sample t-test to calculate the Simpson’s diversity index in R studio.

## Results

We found several dung samples from domestic and wild animal species in the different habitats sampled at BIR. The presence of these different scat samples indicate the variety of domestic animals and wildlife that are present at BIR (Figure 1). The species detected in order of most to least dung counts across all sites were: cattle (2,867), feral pig (127), Rabbit (*Sylvilagu* spp.) (70), white-tailed deer (42), Horse (*Equus ferus caballus)* /Donkey (*Equus asinus)* (11), unknown wildlife (10), Wild Turkey (*Meleagris gallopavo)* (2), Armadillo (*Dasypus novemcinctus)* (1), and Squirrel (*Sciurus spp*.) (1) (Table 2).

We categorized the distribution of dung samples from all sampled block sites in hammock and pasture habitats as dung from a) domestic animals (i.e., cattle, horses, and donkeys), b) feral pigs, and c) all wildlife species combined (Figure 2).

**Figure 2.**
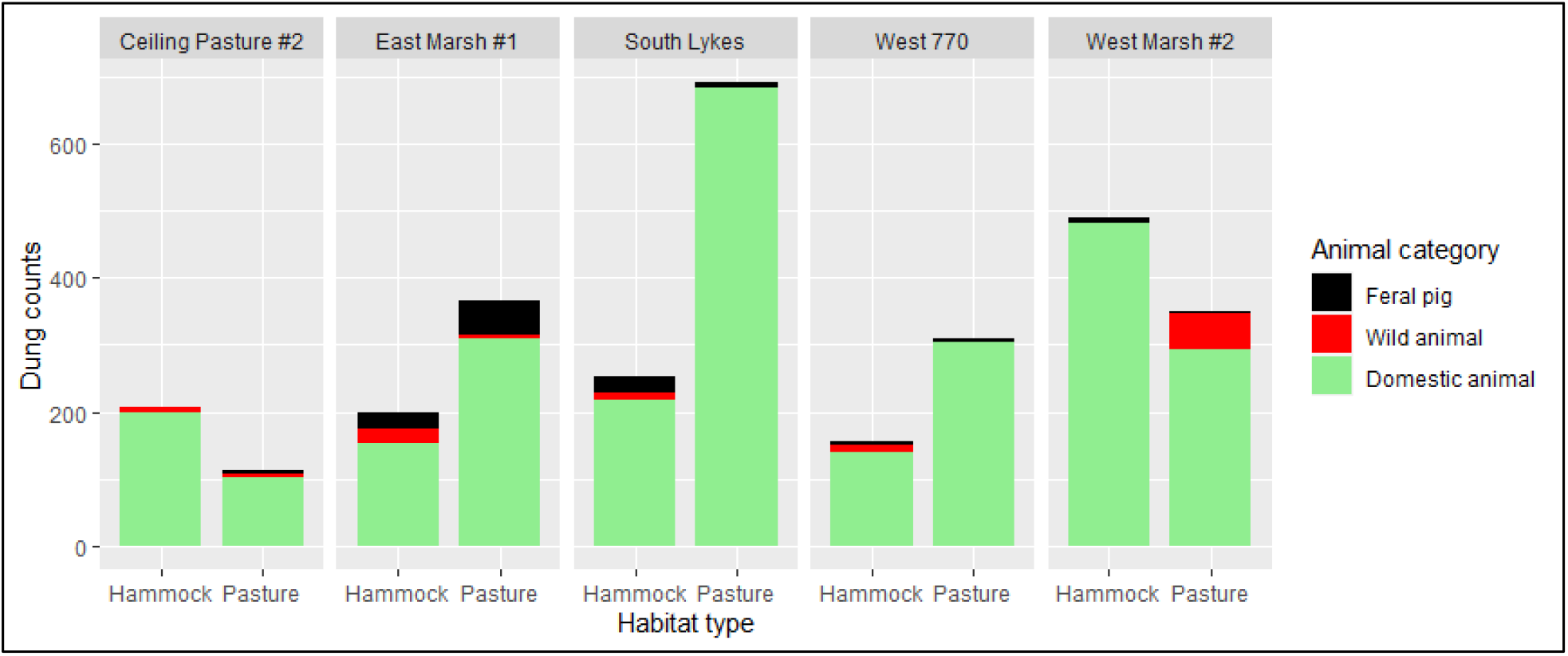
Dung counts of domestic animals, wildlife, and feral pigs observed in various management block sites across hammock and pasture habitats/vegetation types.

*Differences in dung counts* between the hammock and pasture habitats/vegetation type calculated using Wilcoxon signed-rank tests are as follows: difference in dung counts for all animals detected between pasture and hammock(non-significant, p=0.162), differences in cattle dung counts between pasture and hammock (non-significant, p=0.168), differences in feral pig dung counts between pasture and hammock (non-significant, p=0.562), differences in wildlife dung counts between pasture and hammock (non-significant, p=0.134), differences in rabbit dung counts between pasture and hammock (non-significant, p=0.892), differences in deer dung counts between pasture and hammock (non-significant, p=0.066), differences in turkey dung counts between pasture and hammock (non-significant, p=1), differences in armadillo dung counts between pasture and hammock (non-significant, p=1), differences in unknown wildlife dung counts between pasture and hammock (non-significant, p=0.679), and differences in squirrel dung counts between pasture and hammock (non-significant, p=1) (Table 3).

**Table 3.**
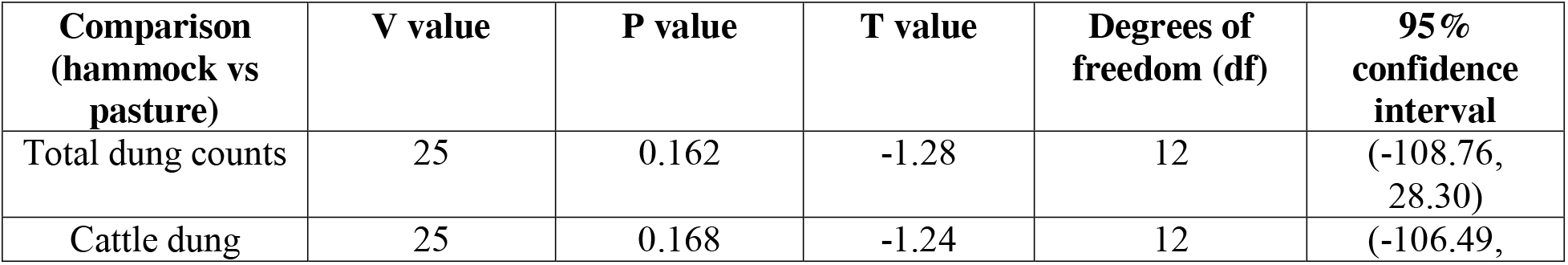
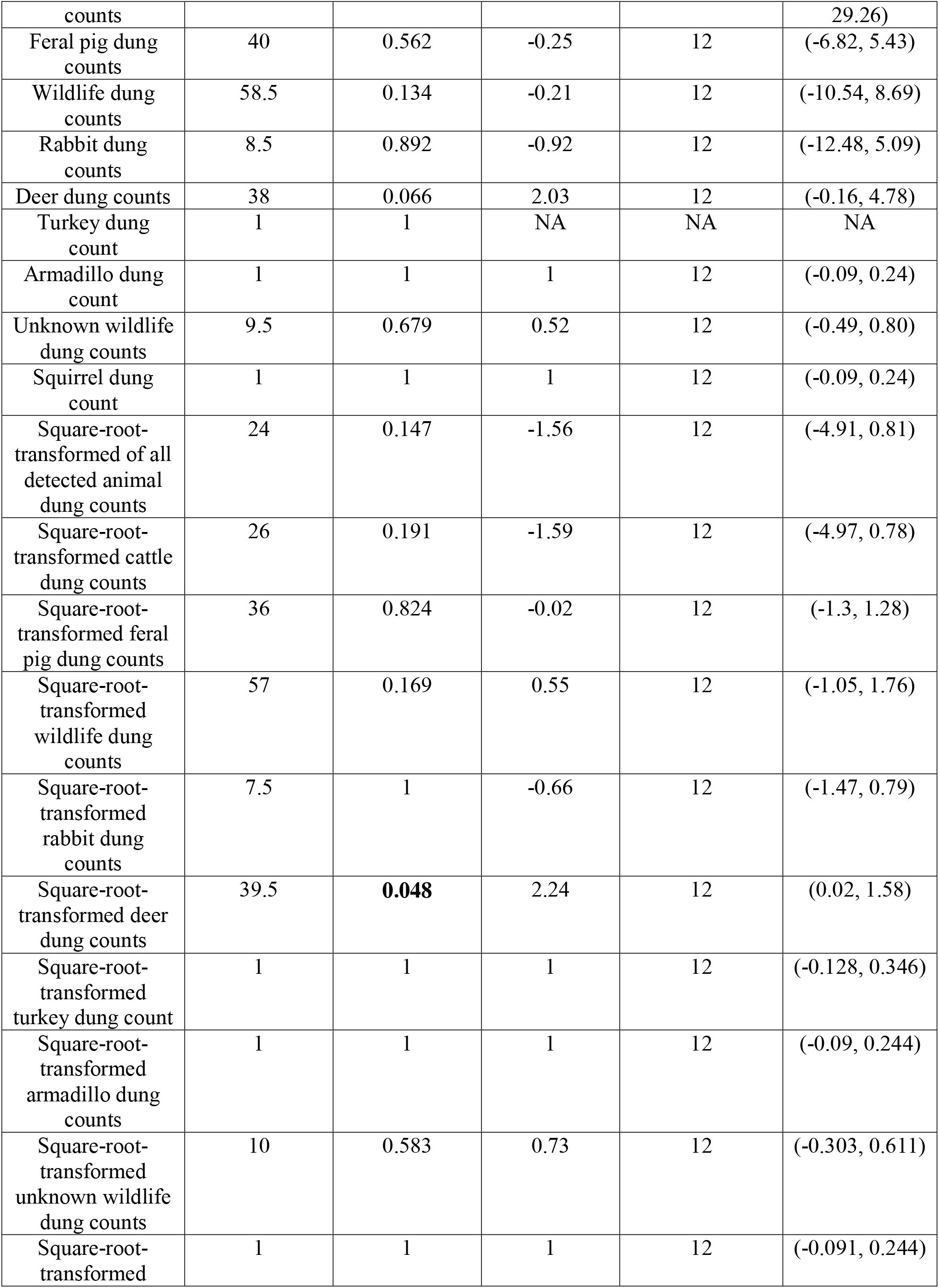
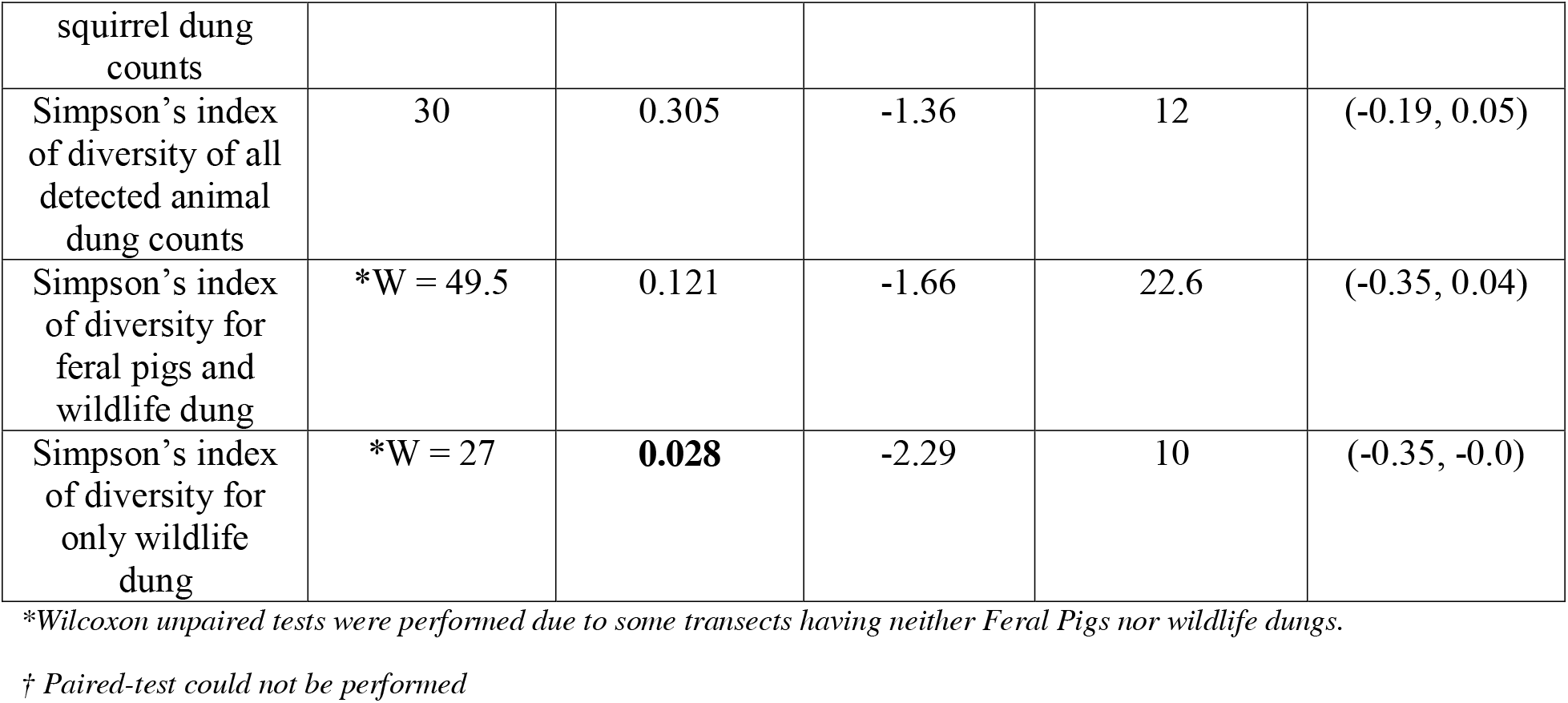
Results of habitat comparisons using Wilcoxon tests and t-tests.

*Differences in square root transformed dung counts* between the hammock and pasture habitats/vegetation type using Wilcoxon signed-rank tests are as follows: differences in dung counts for all animals detected between pasture and hammock (non-significant, p=0.147), differences in cattle dung counts between pasture and hammock (non-significant, p=0.191), differences in feral pig dung counts between pasture and hammock (non-significant, p=0.824), differences in wildlife dung counts between pasture and hammock (non-significant, p=0.169), differences in rabbit dung counts between pasture and hammock (non-significant, p=1.0), differences in deer dung counts between pasture and hammock (**significant, p=0.048**), differences in turkey dung counts between pasture and hammock (non-significant, p=1.0), differences in armadillo dung counts between pasture and hammock (non-significant, p=1.0), differences in unknown wildlife dung counts between pasture and hammock (non-significant, p=0.582), differences in squirrel dung counts between pasture and hammock (non-significant, p=1.0), (Table 3).

The differences in diversity between the habitats calculated using Simpson’s diversity index are as follows: differences in diversity index for all animals detected between pasture and hammock (non-significant, 0.305), differences in diversity index for feral pigs and wildlife between pasture and hammock (non-significant, p=0.121), and differences in diversity index for only wildlife between pasture and hammock (**significant, p=0.028**) (Table 3). Due to small samples sizes, many comparisons were not performed. White-tailed deer dung was detected more often in hammocks than in pastures, and a greater diversity of wildlife dung was observed in hammocks than in pasture habitats. We also observed high counts of cattle dung in both hammocks and pastures suggesting hammocks to be an important area of spatial overlap by wild and domestic animals.

The effect size between cattle dung and feral pig dung was calculated to be Cohen’s d = 2.19 and r = 0.74 whereas between cattle and feral dung samples combined against all wildlife dung was calculated to be Cohen’s d = 1.04 and r = 0.46, indicating large differences in effect sizes between both cattle and feral pig and cattle-feral pig and all other wildlife.

## Discussion

Here, we sought to assess the extent of spatial overlap between wildlife and domestic animals in hammocks and pastures at Buck Island Ranch (BIR), an active cattle ranch in south-central Florida. Based on surveys of animal dung samples, we found that counts of certain wildlife species, such as white-tailed deer, were significantly greater in hammocks than in pastures. We also found that the overall diversity of wildlife species was higher in hammocks than in pastures. Due to the small sample sizes of the observed dung samples, no other comparisons were found to be statistically significant. Cattle dung counts was not found to be significantly different between hammock and pasture habitats, potentially indicating that cattle make considerable use of hammocks. As a result, hammocks appear to be foci of habitat usage for cattle and wildlife.

Another notable finding from this study is that dung presence of cattle and feral pigs were much greater than dung of wildlife as reflected by the dung counts and the effect size calculation. Factors such as body size of animals, predator presence, nutrient availability and quality, defecation frequency, components within scat samples, and local climate are responsible for the large differences in dung samples between cattle and feral pig and cattle-feral pig against all other wildlife that we observed in this study (Hopcraft et al. 2012; Olff et al. 2002; Sinclair et al. 2003; Le Roux et al. 2020). An ongoing question in research on livestock-wildlife interactions is do the abundances of these animal groups trade-off? Interestingly, Yang et al (2021) studied the spatial overlap of feral pigs and cattle at BIR and found that direct feral pig-cattle contacts were occasional, although indirect contacts through resources that are distributed in the ranch, particularly at sites where liquid molasses were kept, were more frequent. In Laikipia, Kenya, Keesing et al (2018) found no direct trade-off in the abundances of wildlife and cattle, but in that study, the abundances of the two animal populations were more equivalent.

As per Kohmann et al (2021), average stocking rate is 152 animal unit days ha^−1^ yr^−1^ in semi-native rangeland and 347 animal unit days’ ha^−1^ yr^−1^ in Pensacola Bahia grass pastures. We could not find estimates for stocking estimates for cattle in hammocks. Even though from the available estimates it seems like cattle might be spending more time in pastures, but from our data, the presence of cattle dung found in large volumes in both pastures and hammocks potentially indicates that cattle are using a range of habitats in BIR. The presence of cattle in pasture and hammock habitats at BIR could impact wildlife movements and presence. The sheer amount of cattle activity in open pastures may also reduce the availability or quality of vegetation, pushing wildlife towards hammocks and other areas which could be more difficult for cattle to access. Hammock habitats may be used over pastures by wildlife for a few distinct reasons. Hammocks provide shade from the hot and humid weather (NPS 2025), they are home to several flora and fauna (Outerbridge 2012) including migratory birds and endangered species (Karim and Main, 2004) and also provide refuge from humans working on the ranch. For wild herbivores, in the hammocks the presence of a diversity of plant species can serve as a source of nutrition (NPS 2021). There were a few wildlife scat samples that we were unable to identify to species, and we identified some of these to be from meso-carnivores; thus, hammocks might also be preferred habitats for hunting by wild animals.

It is important to study how and where wildlife and livestock animals overlap as it can have consequences for transmission of zoonotic and vector-borne diseases (Chakraborty et al. 2023), biodiversity conservation (Lamarque et al. 2009), livestock production (Lamarque et al. 2009), resource competition (Averbeck et al. 2009) and tourism (Genovese et al. 2017). Factors such as local climate, vegetation, food resources, and biological, social, and human activities lead to animal movements (Zengeya et al. 2015). Understanding the mechanisms behind these wildlife-livestock interactions and occurrence of habitat overlap is critical for farmers, ranchers, and land and wildlife managers, as well as crucial in developing effective management programs.

The high volumes of cattle dung presence in both pastures and hammocks at BIR indicate that cattle use of these habitat types might be pushing out some of the wildlife to be concentrated solely in deeper areas of the hammocks. Our study is a snapshot of the spatial overlap occurring between livestock and wildlife at BIR. Our sampling occurred in the summer season in Florida which is characterized by hot and humid weather combined with instances of heavy rainfall. Had this study been performed in a different season, the distribution of cattle or wildlife may have differed. Additionally, scat samples were sometimes dried out due to extended periods of exposure, washed out by rain, or damaged by cattle and vehicle treads, making them difficult to identify. We acknowledge that using scat samples alone comes with many limitations in assessing the relative animal abundance, though in some cases dung counts have been found to be more dependable than visual aerial and ground counts of animals (Barnes 2008, Marques et al. 2001). Lastly, we had limited access to management blocks for data collection since Buck Island Ranch is an active ranch and cattle were grazing continuously in other blocks. Nevertheless, if cattle, feral pigs, and wildlife are regularly present in habitats where there are chances of direct and indirect contact between them, this can be grounds for pathogen spillover (Karmacharya 2024; Chakraborty et al 2023) and depending on livestock management practices can have positive or negative effects.

Apart from pathogen spillover, cattle presence in shared ecosystems can have consequences on the diet and nutritional availability for wild herbivores (Stears and Shrader, 2020) and unintended negative effects on biodiversity (Silori and Mishra 2001). In a review paper by Schieltz and Rubenstein (2016), livestock were found to generally change vegetation structure and cover in ways that can impact small mammals and ungulates. Species adapted to open habitats are often positively affected by grazing, while those that need denser cover for diet or protection against predation, especially small mammals, are negatively affected by livestock grazing. Even though cattle grazing can have both positive and negative effects depending on the season, habitat type, stocking rate, and the geography - in ecosystems shared with wildlife, harmonized coexistence of cattle-wildlife is essential (Barroso and Gortázar 2024).

Therefore, future studies should conduct further analyses of the spatial contacts between cattle and wildlife in different seasons and varying conditions in BIR. This would increase our knowledge on the impacts that cattle grazing and their movements could have on the surrounding wildlife present in BIR. Combining dung counts with other methods such as camera traps and GPS tracking of animals would provide more robust insights into wildlife and cattle interactions.

## Conclusion

In this study, we attempted to determine the spatial overlap between cattle and wildlife on Buck Island Ranch in south-central Florida. We used dung sampling to determine the presence of animals and found that hammock habitats potentially may serve as foci of overlap for wildlife and cattle. Further research should investigate the livestock-wildlife interactions that occur within these habitats to inform conservation, public health, and agricultural measures. Controlling the grazing activities of cattle, creating nutrient-rich hotspots, and having some separation in habitats occupied by cattle and those by wildlife can offset some of the pressures experienced by wildlife (Fynn et al. 2015, Krausman et al. 2009). Additionally, monitoring the areas where cattle-wildlife may interact, and utilizing various grazing systems (such as rotational vs management-intensive), may also reduce the pressures on wildlife as well as help regenerate the vegetation promoting diversity and increased presence of plants, animals, and birds (Fynn et al. 2015, Krausman et al. 2009).

## Acknowledgements

We would like to thank all the research and support staff at Buck Island Ranch, Archbold Biological Station; especially Elizabeth Boughton, Shefali Azad, and Gene Lollis for their tremendous help and support during this project. Funding for this study was obtained through the Graduate College, University of Illinois Dissertation Research Grant. This work was also supported by the USDA National Institute of Food and Agriculture, Hatch project 1026333 (ILLU-875-984).

